# From Genes to Nanotubes: Exploring the UV-Resistome in the Andean Extremophile *Exiguobacterium* Sp. S17

**DOI:** 10.1101/2024.06.28.600890

**Authors:** Fátima Silvina Galván, Daniel Gonzalo Alonso-Reyes, Virginia Helena Albarracín

## Abstract

*Exiguobacterium* sp. S17, a polyextremophile isolated from modern stromatolites in a High-Altitude Andean Lake, exhibits a remarkable multi-resistance profile against toxic arsenic concentrations, high levels of UV radiation (UV), and elevated salinity. Here, we perform a comprehensive characterization of the mechanisms underlying the UV resistance of S17 (UV-resistome/UV_res_) through comparative genomics within the *Exiguobacterium* genus. Additionally, we describe the morphological and ultrastructural changes in the strain when exposed to different levels of UV.

UV_res_ in *Exiguobacterium* species ranges from 112 to 132 genes, with a median of 117. While we anticipated *Exiguobacterium* sp. S17 to lead the non-HAAL UV_res_, it ranked eleventh with 113 genes. This larger UV_res_ in *Exiguobacterium* spp. aligns with their known adaptation to extreme environments. Morphological and ultrastructural analyses using Scanning Electron Microscopy (SEM) and Transmission Electron Microscopy (TEM) demonstrated significant changes in response to UV exposure in S17 cells. We observed the formation of nanotubes (NTs), a novel finding in *Exiguobacterium* spp., which increased with higher UV-B doses. These NTs, confirmed to be membranous structures through sensitivity studies and SEM/TEM imaging, suggest a role in cellular communication and environmental sensing. Genomic evidence supports the presence of essential NT biogenesis genes in S17, further elucidating its adaptive capabilities.

Our study highlights the complex interplay of genetic and phenotypic adaptations enabling *Exiguobacterium* sp. S17 to thrive in extreme UV environments. The novel discovery of NTs under UV stress presents a new avenue for understanding bacterial survival strategies in harsh conditions.

## 1. INTRODUCTION

*Exiguobacterium* is a genus of polyextremophilic bacteria that can survive in a wide range of extreme environmental conditions [1–5]. Indeed, most strains were isolated from acidic and alkaline niches [4, 6], salterns [7], and from ecosystems suffering from extreme temperatures [8–11], high UV radiation (UVR) or high concentrations of heavy metals and metalloids [12–14]. With 17 known species, *Exiguobacterium* has been found to be of interest as a model system for the study of the mechanisms involved in the response to stress and for potential applications in biotechnology, bioremediation, and agriculture [15].

Poly-extremophilic strains of *Exiguobacterium* have been isolated from various Andean microbial ecosystems, including salterns, lakes, and surrounding soils in the Argentinean Puna and Salar de Atacama in Chile, at altitudes ranging from 3,000 to 6,000 meters above sea level [7, 12, 16, 17]. Of particular interest is *Exiguobacterium* sp. S17, the sole strain of this genus isolated from modern stromatolites. S17 was derived from samples taken from the stromatolites along the shores of Lake Socompa (3,570 m, Salta, Argentina). These microbialites inhabit a niche exposed to significant ultraviolet radiation (UVR), with noon UV-B fluxes reaching 10 W m^-2^, alongside high salinity (up to 30%), alkalinity, and arsenic content (32 mg/L), among other physicochemical challenges [18–22].

Previous research has suggested that *Exiguobacterium* sp. S17 exhibits adaptive responses to arsenic and other stressors, facilitating its survival [13, 14]. Described as moderately halotolerant, S17 demonstrates a multi-resistance profile against toxic arsenic concentrations, high ultraviolet radiation (UV) levels, and elevated salinity. Genomic analyses have identified 102 stress response genes in S17, including those related to resistance against various toxic compounds such as antibiotics, arsenic, cadmium, and mercury [14]. Notably, the genome of S17 harbors copies of arsB and acr3 genes, encoding arsenite efflux pumps. Proteomic studies have revealed the overexpression of several genes, including superoxide dismutases, heat shock proteins (such as DnaK chaperone, GroEL, and cpn10), and ABC transporters in response to arsenic exposure [13]. Additionally, it has been demonstrated that S17 exhibits robust cellular aggregation and biofilm formation when exposed to arsenic, suggesting a mechanism for long-term survival under metalloid exposure [23].

Portero et al. (2019) demonstrated S17’s resistance to UV-B radiation and described a UV-resistome (UV_res_), a concept already described and supported by previous studies in Andean microbial extremophiles [24–26]. Ideally, the UV_res_ encompasses various subsystems including UV sensing and response regulators, avoidance and shielding strategies, damage tolerance, oxidative stress response, and energy management, metabolic resetting, and DNA damage repair. Previous investigations have reported several properties and mechanisms of S17 related to these subsystems. For instance, the presence of carotenoid-like compounds in S17 has been documented, suggesting a role in antioxidant defense or photo-protection [26]. Moreover, genes associated with photorepairing enzymes, such as photolyases/cryptochromes (CPF), including a Class I-CPD photolyase, a DASH cryptochrome, and members of the FeS-BCPs subfamily, have been identified [26]. In addition, Ordoñez et al. (2013) [14] reported a complete DNA repair system encoded in the genome, including the UvrABC, and MutL-MutS genes. While some aspects of S17’s UV resistance have been explored, further investigation is needed to elucidate the specific molecular mechanisms involved in adapting to UV radiation in this original environment.

This study aims to comprehensively characterize the mechanisms underlying the UV resistance of *Exiguobacterium* sp. S17. To achieve this, we conducted a comparative bioinformatic analysis of the S17 genome and investigated the morphological and ultrastructural changes in the strain when exposed to varying levels of RUV-B. The findings from this study provide valuable insights into the genetic and cellular mechanisms enabling a poly-extremophile to thrive in environments with high levels of UV.

## 2. MATERIALS AND METHODS

### 2.1. Strains and culture conditions

In the present study, the bacterium *Exiguobacterium* sp. S17, isolated from modern stromatolites from Lake Socompa (3,570 m), located in the Andean Puna in Argentina (S 24°35’34’’ W 68°12’42’’) was used [14, 27]. Bacteria were grown in sterile "H" broth with a pH of 8.0, a halophile-modified Luria-Bertani (LB) medium, at 30°C with moderate agitation at 180 rpm. The medium contained 1% (w/v) tryptone, 0.5% (w/v) yeast extract, 1.5% (w/v) NaCl, 0.3% (w/v) KCl, 0.5% (w/v) MgSO4, and 0.3% (w/v) sodium citrate, with 2% agar added for solid medium when applicable [26]. Bacterial cells were grown to an OD_600nm_ ≈ 0.6 (approximately 10^7^ CFU per milliliter of culture) and immediately used in the designed experiments.

### 2.2. Genome analyses

In this work, we use the previously sequenced and assembled genome of S17 [14]. The assembled genome showed 163 contigs and with a G + C content of 53.14%, and a total of 3,139,227 bases in length. The Whole Genome Shotgun project is available in the database (DB) of NCBI under the accession PRJNA202056. Once the contigs were downloaded from the NCBI Assembly DB, the annotation was implemented using PROKKA [28] with a custom expanded protein DB described by Alonso Reyes & Albarracín (2022). This DB includes Swiss-Prot, TrEMBL, Pfam, SUPERFAMILY, TIGRFAM, and a genus-specific DB built from GenBank files downloaded from the NCBI DB. The same procedure was extended for other 14 genomes from different *Exiguobacterium* strains, whose contigs were also downloaded from NCBI Assembly DB for UV-res comparisons with S17.

The genomes of *Exiguobacterium* sp. S17 and *Exiguobacterium aurantiacum* DSM 6208 (NCBI project PRJNA218043) were queried for the set of genes related to the nanotube assembly using NCBI blast. These genes included the regulator YmdB, hydrolases LytC and LytB, and the CORE proteins FliO, FliP, FliQ, FliR, FlhA, and FlhB. We made a major comparison of the genomic context of S17 and DSM 6208 CORE proteins using the genomic annotations provided by NCBI and the detailed description for *Bacillus subtilis* provided by Bhattacharya et al. (2019).

### 2.3. Effect of UV-B on the cell structure of *Exiguobacterium* sp. S17

To evaluate the effect of ultraviolet radiation (UV-B) on the cell structure of *Exiguobacterium* sp. S17, a quick qualitative assay was performed [30, 31]. S17 strains were grown in H broth at 30°C with shaking (180 rpm) and then harvested in the middle exponential phase (OD_600nm_≈ 0.6), as described in section 2.1. Next, 20 ml aliquots of cell suspension were placed in disposable Petri dishes and exposed to different doses of UV-B irradiance using a Vilbert Lourmat VL-4 lamp, with maximum intensity at 312 nm, for 10 min (0.81 kJ m^-2^), 15 min (1.196 kJ m^-2^), 20 min (1.778 kJ m^-2^), 30 min (2.529 kJ m^-2^), and 40 min (3.268 kJ m^-2^) with shaking at 70 rpm. UV-B irradiance was quantified with a radiometer (Vilbert Lourmat model VLX-3 W) coupled to a UV-B sensor (Vilbert Lourmat model CX-312) under the plastic lid. After UV-B exposure, 1500 μl aliquots were removed from each plate and washed twice with phosphate buffer. The same procedure was performed with control (unexposed) cells. Cells were then fixed with Karnovsky’s fixative for observation by scanning electron microscopy (SEM).

### 2.4. Effect of the membrane detergent sodium dodecyl sulfate (SDS) on nanotube development

To assess whether the nanotubular structures formed by S17 under environmental stress conditions maintain a membranous composition analogous to *Bacillus subtilis* [32], we proposed examining *Exiguobacterium* under varying concentrations of the membrane detergent sodium dodecyl sulfate (SDS). Cells were cultured following the protocol described in section 2.1. A 100 µl sample of the culture was subjected to serial dilutions (10^-1^ to 10^-6^), and 20 µl of each dilution was spread on H agar plates containing SDS concentrations ranging from 0.001% to 0.009%. Microbial growth was assessed after 24 hours of incubation at 30°C. The data obtained were analyzed using the method outlined by [33], where four positive signs (++++), three positive signs (+++), two positive signs (++), one positive sign (+) indicate varying degrees of significant growth, and a negative sign (-) indicates no growth. To visually represent this data, the number of signs corresponded to the count (4, 3, 2, or 1), representing growth units by dilution, which were then summed to derive a single value. Negative signs were treated as zero growth units. Additionally, 100 µl of inoculum was exposed to H broth supplemented with only the maximum (0.009%) and minimum (0.001%) SDS concentrations. This was incubated at 30°C with agitation (180 rpm). Samples were extracted from each test for SEM analysis.

### 2.5. Electron Microscopy

Culture aliquots were collected to examine the morphological and ultrastructural characteristics of strain S17 during the different experiments conducted throughout this work. To do this, the bacterial cells were washed twice in sodium phosphate buffer (pH 7.2). The cells were then fixed with Karnovsky’s fixative (a mixture of 2.66% w/v paraformaldehyde and 1.66% w/v glutaraldehyde in 0.1 M phosphate buffer, pH 7.2), overnight at 4°C. Samples for SEM were prepared following the method described previously [25]. After fixation, cells were washed with 0.1 M phosphate buffer twice and dehydrated in a graded ethanol series (50%, 70%, 90%, and 100%), followed by 100% acetone. Critical point drying was performed using a Denton Vacuum model DCP-1. The samples were then mounted on aluminum supports and subjected to gold sputtering (JEOL model JFC-1100). Finally, the bacterial cells were examined using a Zeiss Supra 55VP scanning electron microscope (Carl Zeiss NTS GmbH, Germany), at the Electron Microscopy Core Facility (CIME-CONICET-UNT).

TEM samples were prepared following the method described previously [25] with slight modifications. After fixation, cells were washed with 0.1 M phosphate buffer twice and embedded in 1.2% agar to form easy-to-handle sample blocks. The samples were post-fixed in 1% osmium tetroxide for 2 hours at 4°C. After washing with phosphate buffer, they were block stained with 2% uranyl acetate for 30 minutes at room temperature in the dark. Subsequently, the agar pellets containing the samples were dehydrated in a graded ethanol series (50%, 70%, 90%, and 100%), followed by 100% acetone. Samples were then infiltrated and embedded in an acetone-SPURR resin sequence (SPURR resin; Ted Pella, Inc.). They were transferred to embedding plates and supplemented with fresh embedding medium. Polymerization was carried out at 60°C for 24 hours. After polymerization, thin sections were cut and mounted on uncoated 200-mesh copper grids (Ted Pella). For negative staining, 10 µl of bacterial culture was placed on Formvar-coated 200 mesh Cu/Pd grids for 5 minutes. Excess liquid was removed using filter paper. Following this, samples were stained with 1% uranyl acetate (Sigma-Aldrich) for 1 minute. The grids were washed for 2 minutes with water and dried. All grids were examined with a transmission electron microscope (Zeiss LIBRA 120; Carl Zeiss AG, Germany) at 80 kV, at the Electron Microscopy Core Facility (CIME-CONICET-UNT).

To examine structural variations in S17 potentially induced by UV exposure, 60 scanning electron micrographs were randomly taken at 5,000× and 10,000× magnifications. A total of 1200 bacteria were analyzed (n=200 per condition). ImageJ 1.54f software (National Institutes of Health, USA) was used for the following measurements: (i) total cell area; and (ii) total number of nanotubes.

### 2.6. Statistical Analysis

Ultrastructural morphometric data, total number of nanotubes, and cell viability analysis based on SDS detergent concentration were compared using one-way ANOVA, followed by Tukey’s post-hoc test (p < 0.05). All statistical and graphical analyses were performed using Prism 10.1.2 software (GraphPad Software, San Diego, CA, USA).

## 3. RESULTS

### 3.1. Genomic features

PROKKA annotation led to 3,330 coding sequences, including 2,630 successfully annotated genes. S17 genomic features were compared to the other fourteen *Exiguobacterium* genomes available in the NCBI assembly database. Supplementary Table S1 shows the annotated genes of all *Exiguobacterium* strains and functions assigned by homology.

### 3.2. UV-resistome of S17

Figure 1 illustrates the number of genes assigned to different subsystems related with the UV_res_ profile for each *Exiguobacterium* spp strain, categorized by types of damage repair, oxidative stress response, and UV avoidance/protection mechanisms. Stacked bars represent the abundance of different genes within each subsystem. While the number of genes varies among strains, a consistent pattern of UV_res_ component abundance is observed across all strains. Notably, S17 ranks eleventh in terms of the number of genes dedicated to the UV_res_, with a total of 113 genes. Unique features of S17’s UV_res_ include a high abundance of genes associated with homologous repair and the presence of an additional copy of the mismatch repair mutS gene (Table S2). Previous work had reported three photolyases in the S17’s genome [26]. Photolyases are very important part of the UV_res_, as they are the finest and most efficient mechanism to repair DNA damage caused by UV-B rays [33–35]. Thus, is important to consider that differences in methodology of photolyase annotation could cause an underestimation of the real capabilities of the UV_res_.

**Figure 1.**
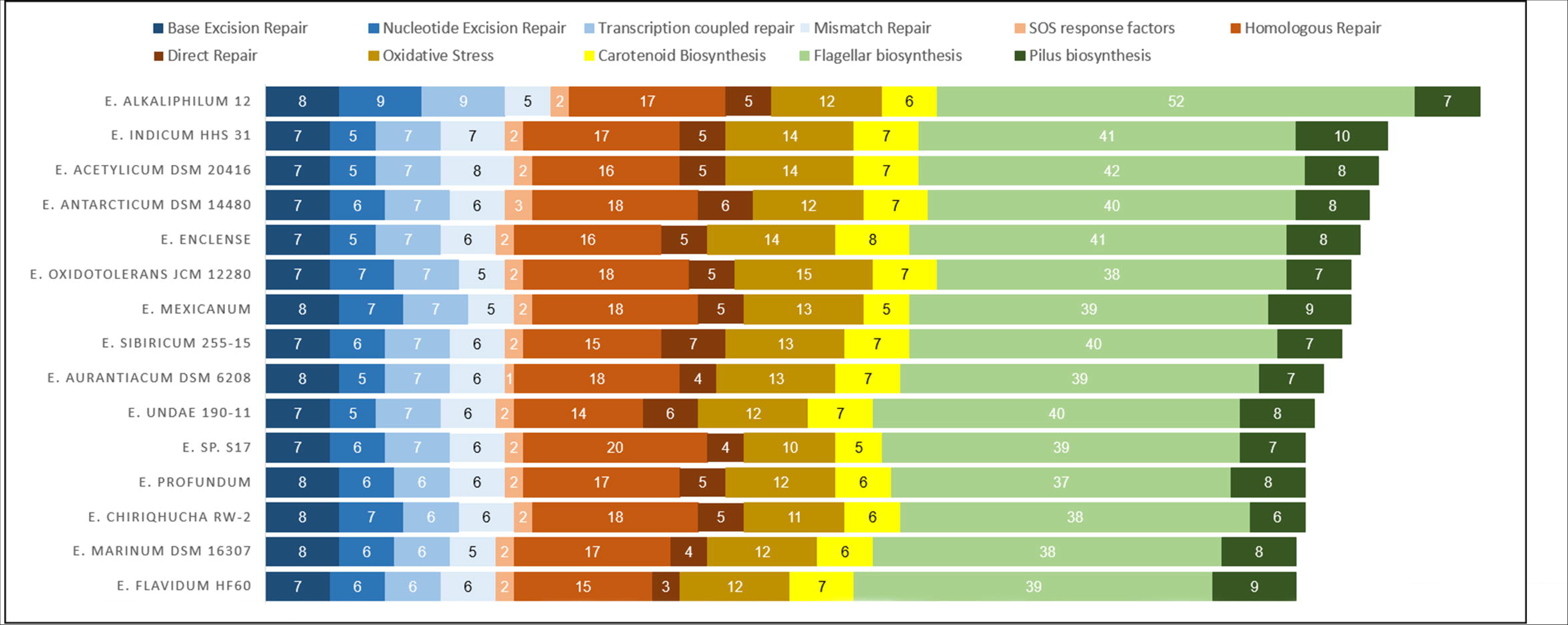
UV-resistomes of *Exiguobacterium* sp. S17 and related strains. The stacked bar chart shows in different colors all the UV-resistome subcategories.

### 3.3. Morphological and ultrastructural analysis of *Exiguobacterium* sp. S17

The cellular morphology of S17 cells was observed under control conditions, not exposed to UV-B radiation. S17 cells typically form coccobacillary aggregates with an average size of 0.8-1.0 µm × 0.6-0.7 µm. These cells maintain a uniform shape with a rough and intact cell surface, often undergoing cell division. SEM and TEM analyses confirmed the preserved and intact cell envelope, displaying the typical structural organization of a Gram-positive bacterium. Extracellular polymeric substances (EPS) were frequently observed surrounding each cell, indicating this strain’s high EPS-producing capacity. Additionally, cytoplasmic structures, including mesosomes and nucleoids, were visualized, highlighting various internal membranous arrangements (Figure 2).

**Figure 2.**
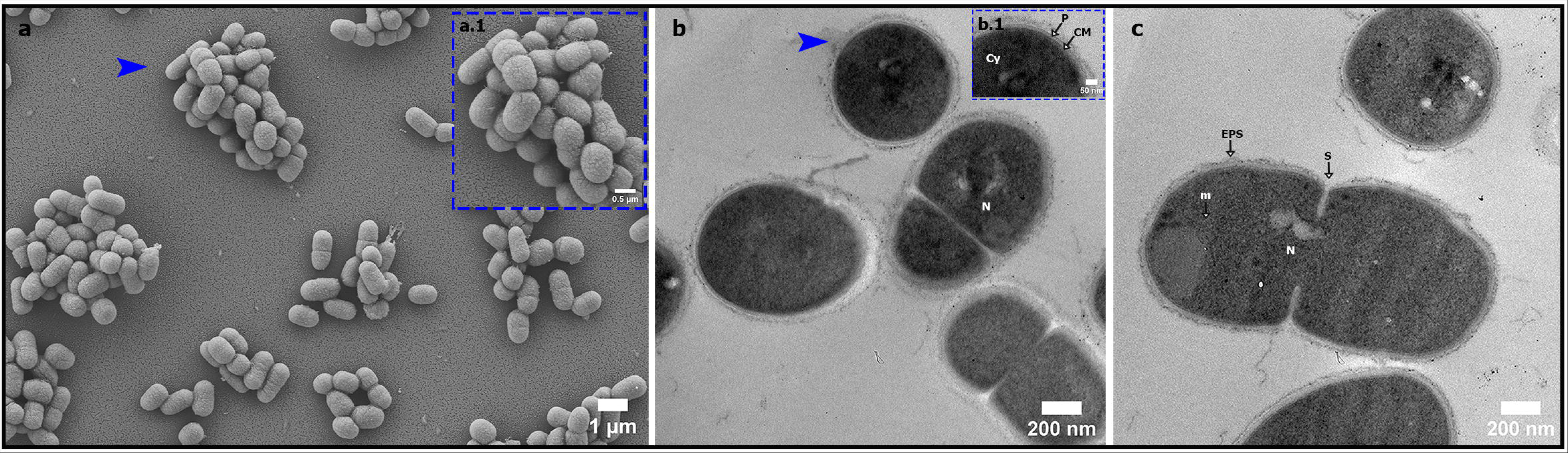
Representative cells of *Exiguobacterium* sp. S17 seen by scanning electron microscopy (SEM) and transmission (TEM). **(a)** Scanning electron micrograph shows cellular aggregates of S17. The scale bar represent 1 um. **(a.1)** A higher magnification image of the region indicated by blue arrow in (a). The scale bar represent 0.5 um. **(b-c)** Transmission electron micrographs show the cross-section of multiple cells, revealing the cytoplasm (Cy), nucleus (N), mesosomes (m), and septal septum (s). The scale bar represents 200 nm. **(b.1)** A higher magnification image of the region indicated by blue arrow in (b) shows the structural organization of the cell wall, with a noticeable thick peptidoglycan layer (P) followed by the cytoplasmic membrane (CM). Typical of Gram-positive bacteria. The scale bar represents 50 nm.

### 3.4. Characterization of S17’s Adaptive Response to UV-B Radiation

To investigate S17’s adaptive response to UV-B radiation, samples were subjected to varying UV exposure doses and analyzed morphological changes using scanning electron microscopy. Following a 20-minute UV exposure, noticeable alterations in cell morphology were observed, although cellular integrity remained largely intact (Figure 3 Left Panel). Intriguingly, increasing UV exposure led to the formation of a dense network of extracellular nanotube-like structures (NTs), connecting cells and exhibiting systematic variations in dimensions (Figure 3 LP [d-f]). NTs were identified by their morphology, as previously described in other bacteria, including the genus *Bacillus* [32, 36, 37]. These membranous structures [32, 38] facilitate the exchange of cellular components (DNA, proteins and metabolites) between neighboring or distant cells [37, 39, 40].

**Figure 3.**
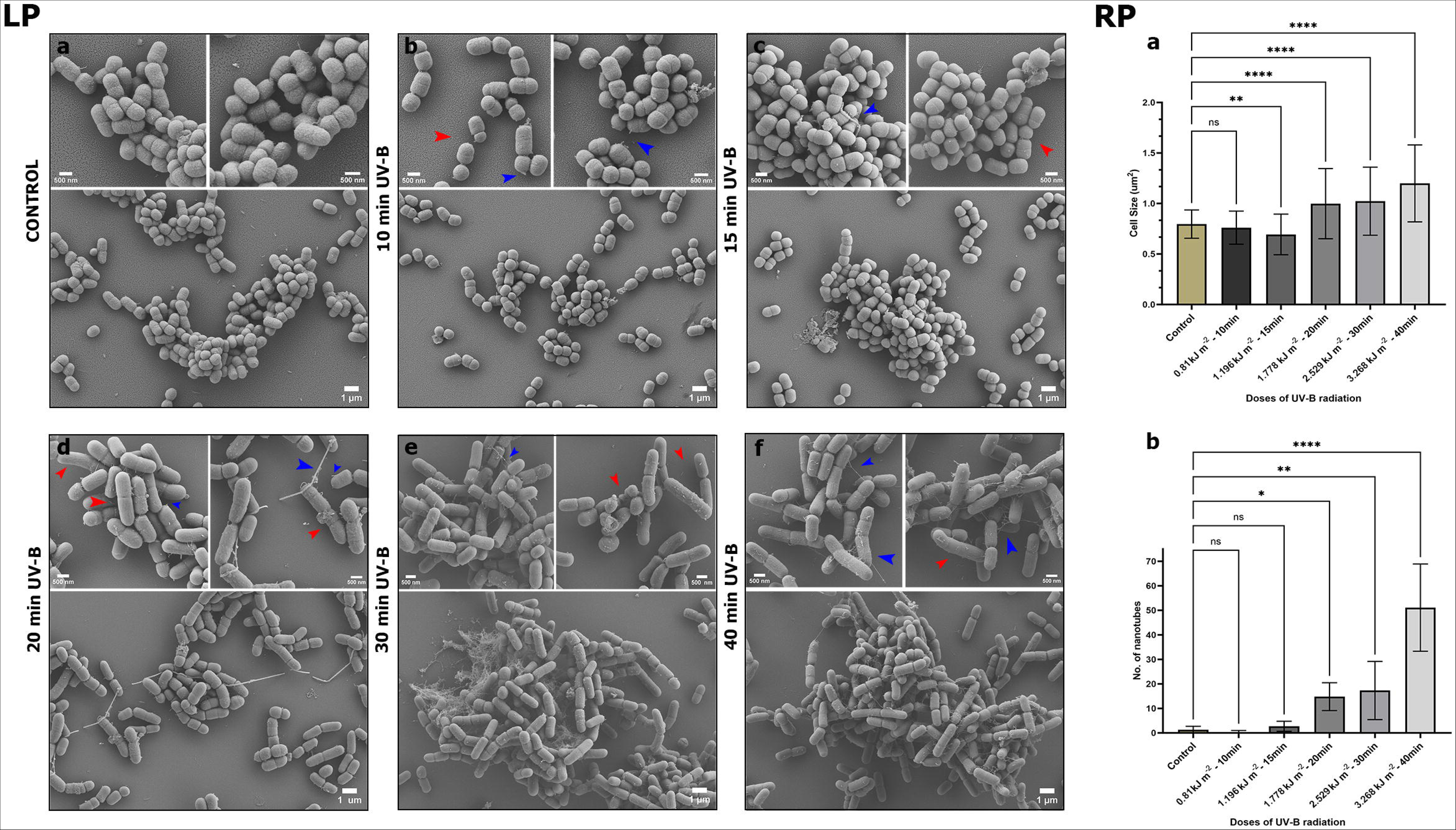
Adaptive response of *Exiguobacterium* sp. S17 to UV-B radiation. **Left Panel (LP)** Scanning electron micrographs show the results of the UV assay. The morphological change due to cell elongation (red arrows) and the production of nanotubes (blue arrows) experienced by the S17 cell over time is perceived. **(a)** Control group **(b)** 10 min, 0.81 kJ m^-2^ **(c)** 15 min, 1.196 kJ m^-2^ **(d)** 20 min, 1.778 kJ m^-2^ **(e)** 30 min, 2.529 kJ m^-2^ **(f)** 40 min, 3.268 kJ m^-2^. Scale bar, 1 μm large boxes, and 500 nm small boxes. **Right Panel (RP)** Quantitative analysis shows a significant increase in cell area **(a)** and nanotube production **(b)** induced by UV radiation. The cell area was 0.9986 ± 0.3479 (mean ± SD) at the dose of 1.778 kJ m^-2^ (20 min), 1.023 ± 0.3372 at the dose of 2.529 kJ m^-2^ (30 min), 1.199 ± 0.3812 at the dose of 3.268 kJ m^-2^ (40 min), compared to the control, 0.7961 ± 0.1402. In the average number of nanotubes, a significant increase was observed at the dose of 3.268 kJ m^-2^ (40 min): 51.10 ± 17.80 (mean ± SD), while the control maintained 1.3 ± 1.418 N° of nanotubes. Asterisks indicate statistical significance determined by one-way ANOVA: **** p < 0.0001, ** p < 0.01, * p < 0.05, and ns: not significant.

### 3.5. Formation of Nanotubes in Response to UV Stress

In S17, NTs exhibited dimensions ranging from 20 to 110 nm in thickness, with thin tubules (diameter 20-40 nm) being the most abundant (Figure 4 LP [a-c]). The nanotubular bridges extended in various directions, covering distances of up to 12 μm. Electron microscopy images demonstrate the various stages of NT formation. Initially, vesicular structures emerge on the cell wall, which then extend outward to form nanotubes. These NTs grow and eventually interconnect, creating a network between cells, and promoting biofilms’ development in many cases (Figure 4 RP [a-e]).

**Figure 4.**
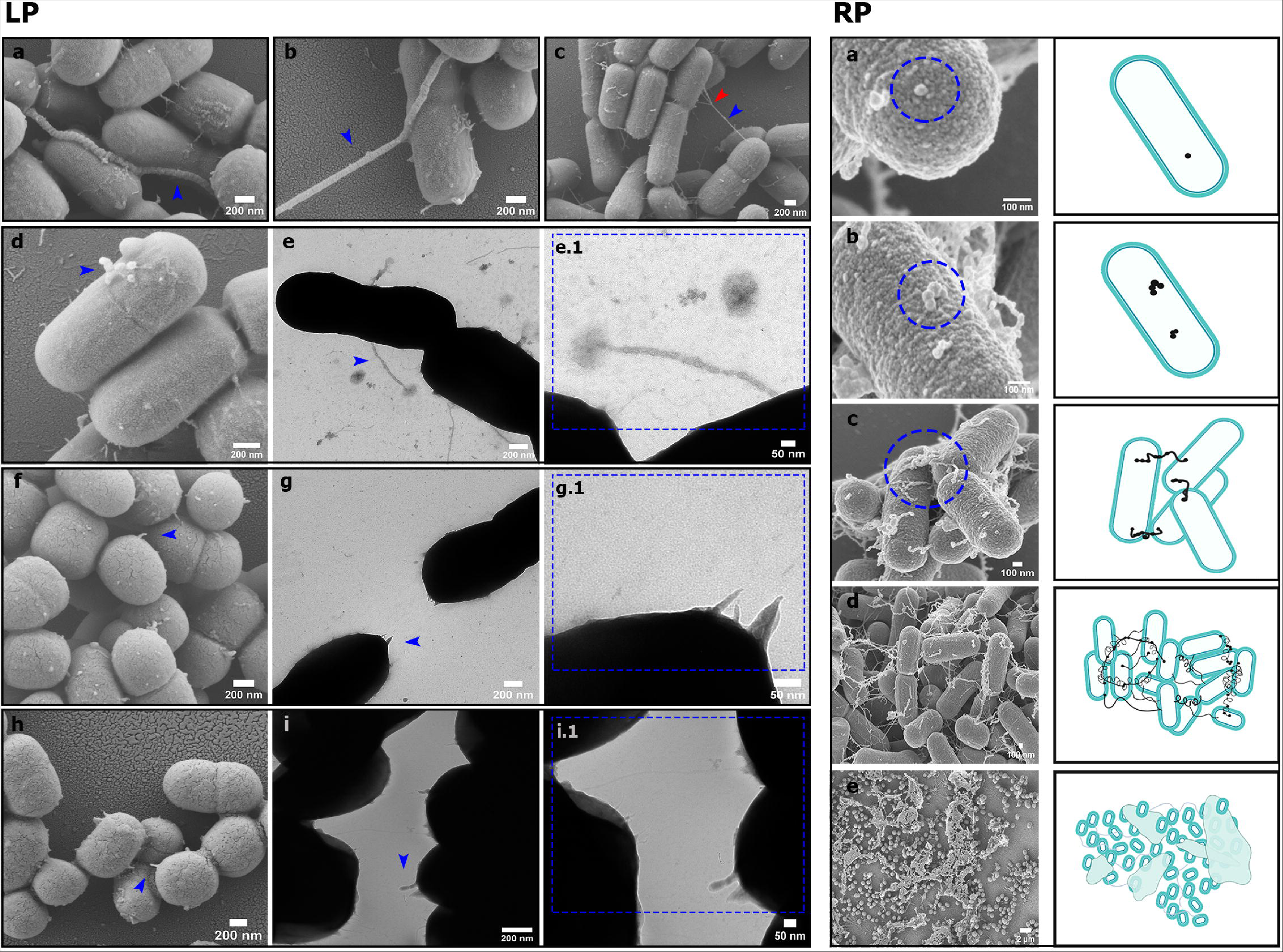
Morphometric variations at the beginning of nanotube formation in S17. **Left Panel (LP)** SEM and TEM (negative stain) images of the tubular protrusions. **(a-c)** Nanotubes of varying dimensions, (a) 95 nm thick and (b) 75 nm thick supercoiled thick tubule (blue arrows), (c) 33 nm thick thin tubule (blue arrow) with intersection node (red Arrow). Scale bars 200 nm. **(d-e)** Correlated SEM/TEM images of the formation of the membranous tubule from nanovesicles adhered to the cell surface (blue arrow). Scale bars 200 nm. **(e.1)** Amplified TEM microphotograph of image (e). Scale bar 50 nm. **(f-g)** Correlated SEM/TEM images of the formation of the nanotube as “arrowhead”-shaped membrane protuberances (blue arrow). Scale bars 200 nm. **(g.1)** Amplified TEM microphotograph of the image (g). Scale bar 50 nm. **(h-i)** Correlated SEM/TEM images of the birth of the nanotube from membrane dilations, acquiring the shape of a drop or tear (blue arrow). Scale bars 200 nm. **(i.1)** Amplified TEM microphotograph of image (i). Scale bar 50 nm. **Right Panel (RP) (a-e)** SEM images and their respective graphic image depict the potential formation of nanotubes from nanovesicles attached to the cell surface. Microphotographs scale bars 100 nm.

The appearance of the NTs was diverse and easily distinguished under higher microscopic magnifications. Various initial formations were observed originating from the bacterial cell membrane, which facilitated the growth of these NTs. Some NTs were seen to emerge from a nanovesicle attached to the cell surface, aligning one after another like a chain, creating constriction points in some regions. Other NTs appeared more uniformly thick (Figure 4 LP [d-e])), a morphological characteristic also extensively documented in *Bacillus subtilis* [38]. Sometimes NTs appear as “arrowhead”-shaped membrane protrusions, which extend consistently and uniformly from the cells to the adjacent bacteria (Figure 4 LP [f-g]). These projections were studied in detail in *Halobacillus* sp. GSS1 [36]. Other ultrastructural variants consisted of NTs with dilations at their end in the shape of a drop or tear (Figure 4 LP [h-i]), a formation that was observed similarly in research carried out with *Flavobacterium anhuiense* [41]. Quantitative analysis revealed a significant increase in both cell size and NT production with higher UV doses (Figure 3 RP [a-b]). The average cell size experienced a notable increase (**** p < 0.0001) after 20 min of continuous UV-B irradiation, showing a 25% increase compared to control cells. After 30 min, the cells were 30% larger while after 40 min their size was 50% larger than that of the average control cell (Figure 3 RP [a]). Considering the density of nanotubes, a notable increase was observed after 20 minutes of UV-B exposure (6-fold higher than unexposed samples). This rise continued with increasing doses, reaching an average of 20 times more nanotubes than the control after 40 minutes of UV-B exposure (Figure 3 RP [b]).

### 3.6. Analysis of the sensitivity of NTs to the membrane detergent sodium dodecyl sulfate (SDS)

To corroborate the effect of the membrane detergent sodium dodecyl sulfate (SDS) on the nanotubes and consequently on the bacterial cell, *Exiguobacterium* sp. S17 was tested against different concentrations of SDS. The sensitivity profile (qualitative analysis) showed that bacterial growth occurred only up to a concentration of 0.003% SDS at the 10^-1^ dilution (Figure 5a). The quantitative analysis reaffirmed this decrease in cell viability, finding a reduction of approximately 20% at a concentration of 0.001% SDS, a 40% reduction in the presence of 0.002% SDS, and a reduction of less than 10% in the presence of 0.003% SDS (Figure 5b). Regarding the nanotubes, SEM analysis of bacterial samples cultured in medium with 0.003% SDS showed minimal development of nanotubes, evidencing the membranous composition that characterizes them [32, 36, 39, 42]. Additionally, cell morphological alterations were observed, potentially due to interruptions in cell division (Figure 5c).

**Figure 5.**
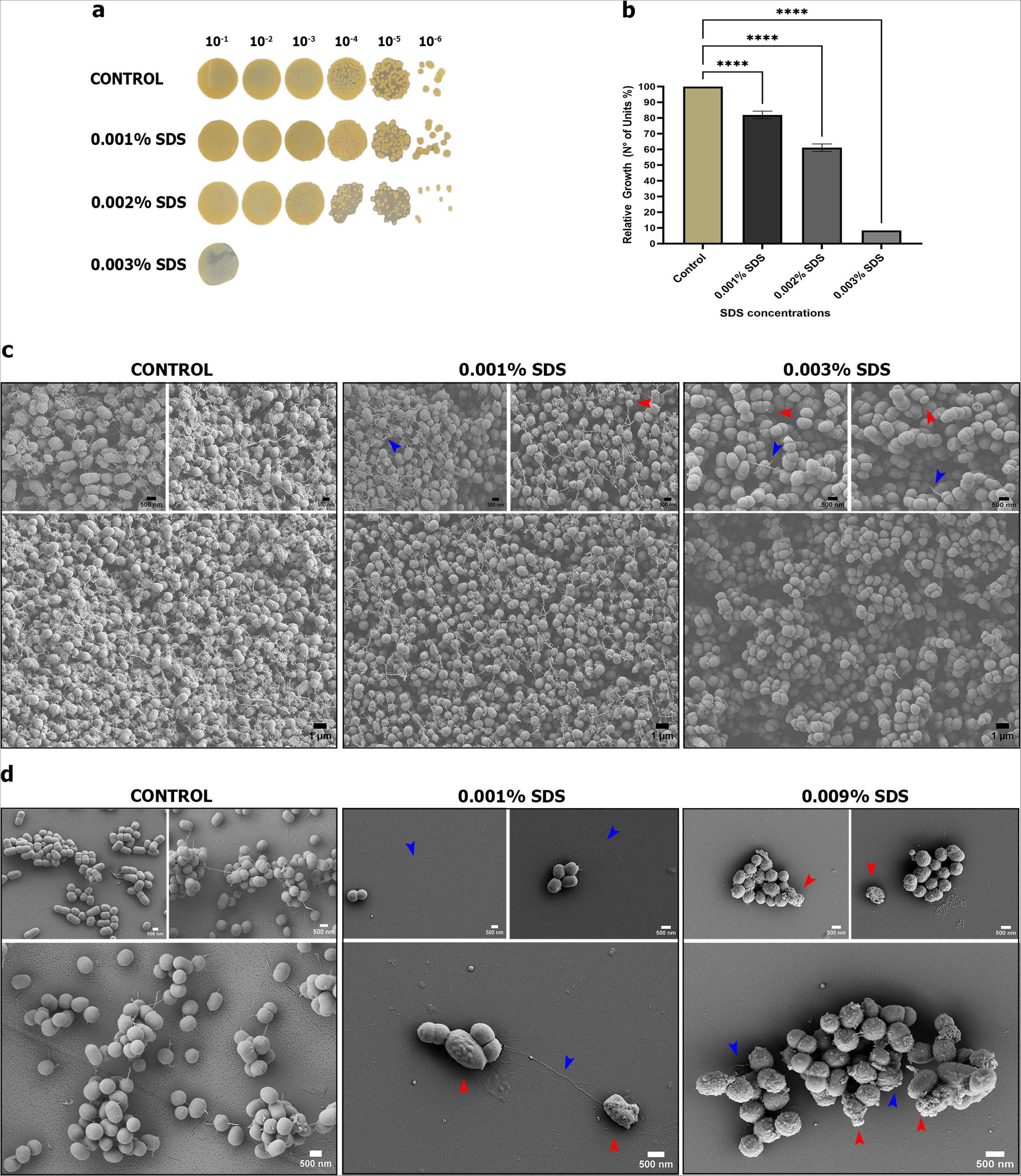
Analysis of the sensitivity of nanotubes to the membrane detergent sodium dodecyl sulfate (SDS). **(a)** Detergent sensitivity analysis of membrane sodium dodecyl sulfate (SDS) in agar medium enriched with increasing concentrations of SDS (0.001% to 0.009%). Only concentrations where colony development occurred (0.001% to 0.003%) are shown. **(b)** Quantitative analyses show that bacterial growth was interrupted with the addition of SDS to the culture medium. The average relative growth percentages with the SDS concentrations of 0.001%, 0.002%, and 0.003% were 81.94 ± 2.40 (No. of units %) (mean ± SD), 61.11 ± 2, 40 (No. of Units %), and 8.33 ± 0.0 (No. of Units %), respectively, compared to the control of 100.0 ± 0.0 (No. of units %). Statistical significance, denoted by asterisks, was determined by one-way ANOVA: **** p < 0.0001. **(c)** SEM micrographs taken from agarized media reveal the absence of nanotube formation at 0.003% SDS concentration (blue arrow) and morphological changes in S17 cells (red arrow). Scale bar, 1 μm large boxes, and 500 nm small boxes. **(d)** SEM micrographs of S17 cells taken from growth in liquid medium spiked with 0.001% SDS and 0.009% SDS. Same solid medium, deterioration in the nanotubular structure and cellular morphological changes with surface variations due to vesicle formation are observed. Scale bar, 500 nm.

S17 cells grown in broth supplemented with 0.001% and 0.009% SDS exhibited minimal growth (OD_600nm_ ≈ 0.4) after 36 hours of incubation. SEM micrographs showed a low number of cellular units under both experimental conditions. The cellular morphological variation, presence of vesicles, and extracellular polymeric material were also highlighted. Damage to NTs was notable, as was their absence in most bacteria (Figure 5d).

### 3.7. Genomic evidence of NTs biogenesis in S17

Through genomic analysis, we identified key genes involved in NT formation, including ymdB, the CORE protein complex, and LytC. These genes are essential for NT production, biogenesis, and cell wall remodeling. The YmdB protein encoded by the *ymdB* gene is the key enzyme for NT biogenesis; it acts as a phosphodiesterase which hydrolyzes cyclic nucleotides like cAMP. It is located throughout the entire NT length, and is critical for intercellular molecular exchange. It also intervenes as a key regulator in decision-making processes for biofilm formation, motility, and sporulation [38, 43–45]. Notably, the presence of these genes in S17’s genome (Table 1) supports the experimental findings of NT formation and confirms S17’s NTs as the first studied in an *Exiguobacterium* strain. Furthermore, *ymdB* gene was found in most of the set of *Exiguobacterium* genomes through a BLAST search (data not shown). The CORE protein complex is composed of five integral membrane proteins (FliP, FliQ, FliR, FlhB, and FlhA) and FliO, a non-structural component that serves as a scaffold for the assembly of CORE. This complex acts as a platform for flagellum and NT biogenesis [29]. In Figure 6, we illustrate the *fla/che operon* of *Exiguobacterium*, indicating the CORE constituent gene locations. Ultrastructural analysis revealed NTs emerging directly from the cell membrane, traversing the wall to protrude from the cell surface. This process is enabled by the expression of the wall hydrolase LytC (N-cetylmuramoyl-L-alanine amidase), which participates in the remodeling of the cell wall, facilitating the extrusion and penetration of NTs into the recipient bacteria [46]. In *Bacillus subtilis*, LytC activity is enhanced in the presence of LytB, an activator. However, our bioinformatics research did not identify the genetic sequence for LytB in S17, which indicates the recruitment of other hydrolases in conjunction with LytC to aid NT penetration into recipient bacteria.

**Figure 6.**
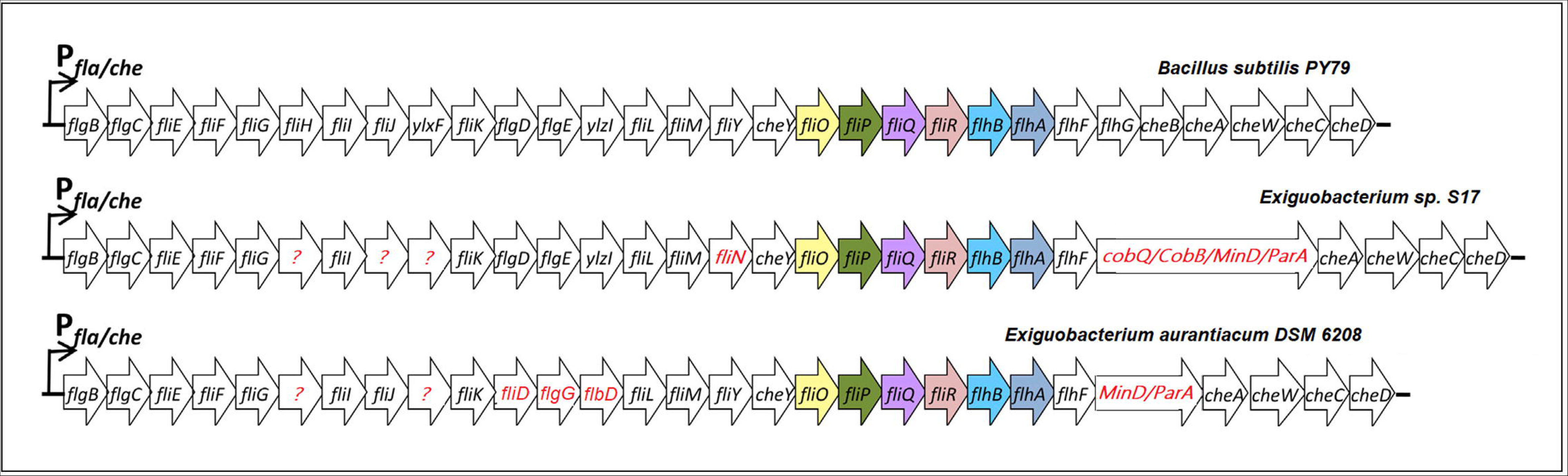
Schematic figure representing the fla/che operon for *B. subtilis* PY79 (top), *E.* sp. S17 (center) and *E. aurantiacum* DSM 6208 (bottom). The position of the genes that encode the protein components necessary for the formation of the flagellar basal body, the CORE complex, is presented. Differences in *Bacillus* patterns, observed in *Exiguobacterium* operons annotated by the NCBI PGAP, are written in red.

**Table 1.** Report of genes required for nanotube assembly in *Exiguobacterium* sp. S17 and *Exiguobacterium aurantiacum* DSM 6208.

| Gene | Function | *Exiguobacterium*<br>sp. S17 | *Exiguobacterium aurantiacum*<br>DSM 6208 |
| --- | --- | --- | --- |
| YmdB | Required for<br>nanotube<br>production | Present | Present |
| LytC | Facilitate<br>nanotube<br>penetration in<br>the cell wall. | Present | Present |
| LytB |  | Absent | Absent |
| FliO | Nanotube<br>assembly<br>(CORE) | Present | Present |
| FliP |  | Present | Present |
| FliQ |  | Present | Present |
| FliR |  | Present | Present |
| FlhA |  | Present | Present |
| FlhB |  | Present | Present |

## 4. DISCUSSION

The high UV resistance observed in S17 prompts a thorough investigation into its underlying mechanisms, highlighting integrated physiological and molecular responses triggered by ultraviolet light exposure. We have termed this system the "UV-resistome," as previously described for other extremophilic microorganisms [24–26]. Essentially, the UV_res_ relies on the expression of a diverse array of genes aimed at either evading or repairing UV-induced damage. These genes integrate into several subsystems, including UV sensing and response regulators, UV-avoidance and shielding strategies, damage tolerance and oxidative stress response, energy management and metabolic resetting, and DNA damage repair. Therefore, we conducted a comprehensive screening of genes associated with each UV_res_ subsystem for fifteen *Exiguobacterium* genomes, including strain S17. This analysis unveiled the relative genomic potential of S17 in defending against UV-B radiation, as it naturally endures the highest irradiation levels on Earth. Similar to the approach by Alonso-Reyes et al. (2021), we explored UV resistance holistically, incorporating genes that may contribute to UV evasion responses or mitigate its negative impacts, such as motility genes, pilus formation, and gas vesicle synthesis [47].

Previous genomic and metagenomic studies have demonstrated a relationship between UV resistance profile and the number of specific genes, indicating that higher the genes involved in the UV_res_ the better adaptation to high UV irradiance environments [25, 48, 49]. High-Altitude Andean Lakes (HAAL) strains and communities exhibit exceptional genetic configurations, which provide them with superior resilience to UV radiation compared to non-HAAL counterparts. For instance, the HAAL strain *Nesterenkonia* sp. Act20 possesses the largest UV_res_ gene count within its genus, ranging from 56 to 114 genes, with a median of 78, positioning Act20 at the top with 114 genes [25]. In the current study, we observe that the UV_res_ in *Exiguobacterium* species ranges from 112 to 132 genes, with a median of 117. While we anticipated *Exiguobacterium* sp. S17 to lead the non-HAAL UV_res_, it ranked eleventh with 113 genes, slightly below the median but still comparable to Act20. This larger UV_res_ in *Exiguobacterium* spp. aligns with their known adaptation to extreme environments.

Experimental data highlight S17’s resilience; agar tests showed its UV-B resistance comparable to *Exiguobacterium aurantiacum* DSM 6208 after 30 minutes of UV exposure but significantly stronger after 60 minutes [26]. Despite this, both strains have similar UV_res_ gene counts (113 for S17 and 115 for DSM 6208; see Table S2). This suggests additional factors beyond UV_res_ gene count, such as gene expression, contribute to S17’s enhanced long-term resistance. Future transcriptomic and proteomic studies should investigate these factors to explain S17’s superior adaptation to HAAL environments.

Previously, we reported that Act20’s UV_res_ includes genes for ectoine synthesis, a photoprotective pigment, along with genes for swarming motility and gas vesicle synthesis, aiding UV evasion/shielding. In contrast, S17 lacks these traits (Table S2). Instead, S17’s genome includes genes for pili-associated motility (twitching motility), absent in Act20, and a more extensive set of flagellar genes (39 in S17 vs. 22 in Act20). S17 also has a comprehensive set of genes for Homologous Repair (HR), which is crucial for repairing double-strand breaks (DSBs) and other UV-induced lesions like CPDs and (6-4)-PPs. HR repair, potentially more critical than Nucleotide Excision Repair (NER) and Base Excision Repair (BER) for UV damage, can cause genome rearrangements and mutations, promoting adaptation in highly UV-exposed environments [50] [51].

Beyond these molecular repair mechanisms, bacteria employ several survival strategies under adverse conditions, including the synthesis of secondary metabolites, biofilm formation, sporulation, and conjugation [52]. Our findings reveal that the adaption to UV of HAAĹs bacteria can also depend on the cellular communication via nanotubes. These membranous tubes connect cells, facilitate the exchange of cellular components, and play a critical role in environmental sensing. The formation of multiple nanotubular protrusions was documented in *Halobacillus* sp. GSS1, a halophile from the Sundarbans mangrove, in response to nutritional stress, where cell-cell bonding occurred mainly between living and dead cells, indicating a possible utilization of cellular debris as nutrients by intact cells [36]. In contrast, *Pospíšil* et al. (2020) observed that NTs in *B. subtilis* were associated with stress conditions such as biophysical forces or the presence of antibiotics, highlighting their formation in dying cells, eradicating the possibility of molecular transfer through the nanotubular conduit. However, the claim about the production of NTs from living organisms and their role in intercellular communication gains support in current research conducted on marine bacteria and cyanobacteria: *Alteromonas* sp. ALTSIO and *Pseudoalteromonas* sp. TW7, *Synechococcus* sp. and *Prochlorococcus* sp., respectively. These microorganisms promoted the development of direct cellular connections through tubules of variable diameters due to the extensive oligotrophic zones they inhabit, which facilitates the efficient management of nutrients within their communities [53, 54]. In this context, it is worth mentioning that while nanotube development has been reported in several bacterial species, this study represents the first report documenting observations of intercellular nanotubes in the extremophilic strain *Exiguobacterium* using high-resolution TEM and SEM microscopy. Furthermore, an increase in NT formation was evidenced in response to high doses of UV-B, a phenomenon not previously documented in bacteria [52][38].

SEM and TEM analyses, alongside sensitivity studies using SDS, confirmed the membranous composition and multilayer arrangement of NTs, indicating lipid bilayer continuity. We also elucidated the various stages of NT genesis: membranous tubes initially forming from nascent vesicles, subsequently elongating and producing regular constrictions (pearl tubes), and ultimately resulting in a network of interconnected tubes of varying diameters [32]. A similar mechanisms on NTs biogenesis was previously described for *Bacillus subtilis* NTs using Cryo-EM [38].

In accordance, S17 genome revealed all the essential building blocks for NT biogenesis: i) YmdB protein, crucial for NT production and cytoplasmic compound transport, ii) CORE protein complex, serving as a platform for NT biogenesis and ejection, and iii) cell wall remodeling enzyme LytC [24, 25]. This genomic evidence supports our experimental findings, confirming the presence and unique formation of NTs in S17, marking the first detailed study of NTs in *Exiguobacterium* strains. Intriguingly, these genes were absent in other HAAL genomes of UV-resistant strains such as *Acinetobacter* sp. Ver3 and *Nesterenkonia* sp. Act20.

## 5. CONCLUSION

This study has provided significant insights into the UV-resistome of *Exiguobacterium* sp. S17, a polyextremophile isolated from modern stromatolites in a high-altitude Andean lake. Through comprehensive genomic and morphological analyses, we have identified key genes and cellular adaptations that contribute to this strain’s remarkable resistance to UV radiation. Notably, the discovery of nanotube formation under UV stress highlights a novel aspect of bacterial adaptation, suggesting a potential mechanism for cellular communication and environmental sensing.

The identification of 113 genes related to the UV-resistome in S17, including multiple photolyases and homologous repair genes, underscores the complexity of its UV resistance strategies. The presence of nanotubes and their increase in response to higher UV-B doses suggest an adaptive response that warrants further investigation. These findings contribute to our understanding of bacterial survival mechanisms in extreme environments and open new avenues for biotechnological applications.

Future research will focus on functional studies of the identified UV-resistome genes to understand their specific roles in UV resistance, as well as investigate the role of nanotubes in cellular communication and stress response. Comparative genomic analyses involving diverse extremophiles are underway to identify common and unique UV resistance strategies. Additionally, proteomic and metabolomic profiling under different UV conditions are being conducted to provide insights into changes in protein expression and metabolic pathways.

## ACKNOWLEDGEMENTS

The authors acknowledge the generous financial support by PICT 2019-3216 projects. VHA is a researcher from the National Research Council (CONICET) in Argentina. FSG and DGA-R are the recipients of doctoral and post-doctoral fellowships, respectively, from CONICET. Electron micrographs used in this study were taken at the Center for Electron Microscopy (CIME) belonging to UNT and CONICET, in Tucumán, Argentina. This manuscript has been released as a Pre-Print at bioRxiv.

